# The S1PR modulator Ozanimod reverses spine deficits, Amyloid-β plaques deposition and neuroinflammation in aged APP/PS1 mice

**DOI:** 10.64898/2026.05.02.722389

**Authors:** Rieke Fritz, Thomas Endres, Lukas Schönwolf, Georgia-Ioanna Kartalou, Kurt Gottmann, Volkmar Leßmann

## Abstract

Innovative therapeutic approaches providing clinically effective medication for Alzheimer’s disease (AD) patients are urgently needed. In recent years, monoclonal antibodies against Amyloid-β (Aβ) oligomers were approved as the first disease modifying AD therapies. However, their effects on cognitive decline of AD patients are still limited – most likely because neuroinflammatory processes downstream of Aβ plaques remain activated, even when plaques are depleted. Accordingly, anti-inflammatory drugs are currently considered as valuable (co-)treatments to target Aβ associated neuro-inflammation. The anti-inflammatory sphingosine-1-phosphate receptor (S1PR) modulator fingolimod (FTY720) has been shown to alleviate synaptic and cognitive deficits in numerous mouse models of AD. Whether other S1PR modulators exert similar beneficial effects is largely unknown. Here we used a transgenic APP/PS1 AD mouse model to investigate whether the S1PR modulator Ozanimod (RPC1063) can rescue AD pathology and synaptic dysfunctions even when treatment is initiated at 16-17 months of age, which is 10 months after onset of cognitive deficits. We performed quantitative dendritic spine analysis in hippocampal CA1 pyramidal neurons and immunohistochemical labelling of Iba1-positive activated microglia, GFAP-positive reactive astrocytes, and Aβ plaques in hippocampus and neocortex. Our results reveal that 6 weeks of Ozanimod treatment via drinking water rescues synaptic spine deficits, counteracts Aβ pathology and reduces neuro-inflammation in the hippocampus and neocortex of APP/PS1 mice. Therefore, it might hold promise to examine a potential disease-modifying effect of S1PR modulators in clinical trials with AD patients.

## Introduction

Alzheimer’s disease and related forms of dementia are major threats for healthy aging in most post-industrial countries. Since the incidence of dementia drastically increases beyond 65 years of age, the welcome increased longevity in these countries drives costs for health services and care for demented patients to unaffordable heights. Currently, more than 50 million people worldwide are living with dementia, and this number is projected to triple by 2050 (Livingston *et al*., 2017; Livingston *et al*., 2024). Consequently, developing a pharmacological treatment that effectively alleviates cognitive deficits in demented patients is critically needed.

Alzheimer’s disease (AD) is the most common (∼60%) form of dementia and is thought to be caused by formation of extracellular Amyloid-β (Aβ) plaques, starting in neocortex ∼15 years before onset of cognitive symptoms, followed by formation of intracellular tau-positive neurofibrillary tangles (NFTs) that are first observed in trans-entorhinal cortex and hippocampus (Jack *et al*., 2016; Congdon & Sigurdsson, 2018; Jansen *et al*., 2019). This primary pathology gives rise to neuroinflammatory responses mediated by microglia (Qin *et al*., 2017; Salter & Stevens, 2017; Paolicelli *et al*., 2022), reactive astrocytes and possibly T-lymphocytes that trigger synaptic deficits and neural circuit dysfunctions, leading to progressive cognitive decline and loss of memory in affected patients.

Starting in 2013, the sphingosine-1-phosphate (S1P) receptor modulator Fingolimod (FTY720) was reported to protect neurons from Aβ-toxicity ((Asle-Rousta *et al*., 2013; Doi *et al*., 2013; Hemmati *et al*., 2013; Fukumoto *et al*., 2014; Ruiz *et al*., 2014)) reviewed e.g. in ((Angelopoulou & Piperi, 2019); (Bascunana *et al*., 2020)), but its cellular mode of action remained unclear. More recently, anti-inflammatory treatment with fingolimod has emerged to mitigate synaptic and memory deficits, and to reduce Aβ- and tau-pathology in a wide range of transgenic AD mouse models (Aytan *et al*., 2016) (Kartalou *et al*., 2020b; Baloni *et al*., 2022; Fagan *et al*., 2022) reviewed e.g. in (Pournajaf *et al*., 2022; Lessmann *et al*., 2023). Fingolimod tackles the detrimental microglia- and astrocyte-mediated immunological and neuroinflammatory processes that are triggered by amyloid-β and tau fibrillary tangle pathology, even when it is applied after disease onset (Kartalou *et al*., 2020b; Baloni *et al*., 2022; Fagan *et al*., 2022; Bascunana *et al*., 2023). Such a medication that is effective even after onset of memory dysfunctions is crucial to allow successful treatment of human patients who are already diagnosed with AD.

In target tissue, Fingolimod is converted by sphingosine kinase 2 (SphK2) to Fingolimod-phosphate (an analog of endogenous S1P), which binds with comparable affinity to all five subtypes of G-protein coupled S1P receptors (S1PR1-S1PR5) that are widely expressed across mammalian tissue (Angelopoulou & Piperi, 2019). S1PRs are found in numerous hematopoietic and immune cells, including microglia and t-lymphocytes but are also involved in modulating heart and vascular functions (Chaudhry *et al*., 2017; Comi *et al*., 2017; Roy *et al*., 2021; Mirzaei *et al*., 2022). Recent studies suggest a role of fingolimod signaling in suppressing intracellular tau-phosphorylation (Wang et al., 2021; Yin et al., 2021), which represents a hallmark of AD pathology in human patients.

A genome wide association study in human patients (Baloni *et al*., 2022) provided strong evidence that mutations in genes encoding enzymes of the sphingosine/ceramide (SM) pathway are associated with an increased risk to develop sporadic AD. This study identified genetic variants in seven of the 35 genes in the SM pathway to be significantly associated with AD and its (bio)markers, covering the whole spectrum of Amyloid-β, Tau, Neurodegeneration, Cognition (A-T-N-C) measures of AD (Jack *et al*., 2018). Since sphingosine-1-phospate (S1P) – the natural ligand of S1P receptors – is a reaction product of the SM pathway, and Fingolimod-phosphate mimics S1P effects, the SM pathway is an obvious drug target for AD treatment. This is consistent with the observation that reduced S1P and S1PR-1 levels in the brain of post mortem AD patients negatively correlate with Aβ plaque burden and disease severity (Ceccom *et al*., 2014a; Ceccom *et al*., 2014b; Couttas *et al*., 2014). As fingolimod-phosphate (i.e., the active form of FTY720 in the brain) compensates reduced S1P levels, it can counteract decreased S1PR signaling in AD patients.

Fingolimod was approved in 2010 for the treatment of Multiple Sclerosis. Meanwhile, several S1PR subtype-specific analogs of Fingolimod (e.g. Ozanimod, Siponimod, Ponesimod; all approved for treatment of relapsing or progressive MS by FDA and EMA (Comi *et al*., 2019; Benedict *et al*., 2021; Kappos *et al*., 2021; Sandborn *et al*., 2021)) were developed to reduce unwanted side effects of Fingolimod such as bradycardia and the required phosphorylation for activation of the drug. However, whether these substances have the same beneficial effects against AD pathology and synaptic deficits is currently unclear (but see (Zhu *et al*., 2023)).

In the current study, we tested whether one month of oral application of Ozanimod via drinking water to APP/PS1 AD mice, starting at 16 months of age, can reduce Aβ-pathology, synaptic deficits, and microglia/astroglia-mediated neuroinflammation. Our results demonstrate a similar beneficial effect of Ozanimod against AD pathology as previously observed with fingolimod. Given the lack of bradycardia side effects, this identifies Ozanimod as a promising S1PR modulator that might provide an effective treatment of AD patients, especially for those suffering cardiac co-morbidities.

## Results

### 1. Effects of Ozanimod on spine pathology in APP/PS1 mice

In the APP/PS1 mouse model we use (Radde *et al*., 2006), AD plaque formation has started to develop at 3 months of age in hippocampus and neocortex, whereas synaptic dysfunctions, including spine reduction, impaired LTP, and deficits in hippocampus-dependent spatial and fear learning can be detected around 6 months of age ((Kartalou *et al*., 2020b); compare **Fig.1**). We therefore wanted to determine whether start of Ozanimod treatment 10 months after established cognitive and synaptic deficits can still successfully counteract spine pathology in APP/PS1 mice. To this aim, 16-17 months old heterozygous APP/ PS1 mice received 1 mg/kg/day Ozanimod for 45 days via the drinking water (compare **Fig.1**). As we have shown previously that synaptic pathology in this AD mouse model can be reliably quantified by Golgi Cox-staining of dendritic spines in secondary dendrites of hippocampal CA1 pyramidal cells (Kartalou *et al*., 2020a; Kartalou *et al*., 2020b), we used this method to analyze synaptic deficits.

**Figure 1.**
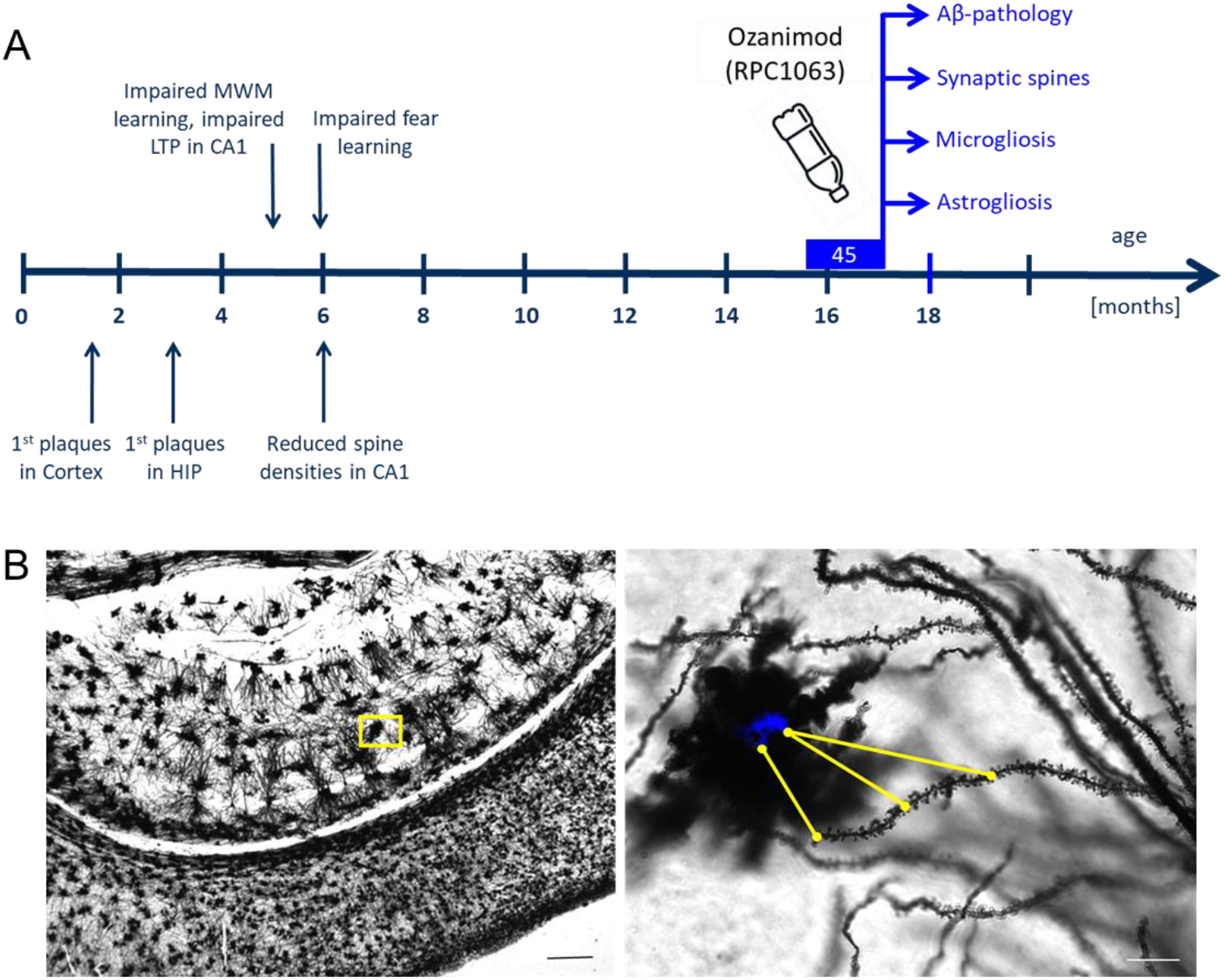
Timeline of APP/PS1 mouse deficits, Ozanimod treatment regime and read-out of results. **A)** Ozanimod was administered to heterozygous APP/PS1 mice via drinking water at a concentration of 1 mg/kg bodyweight per day. Ozanimod application started at 16 months, when fully developed memory impairments, synaptic deficits, and neuroinflammation have persisted for 10 months without treatment. Ozanimod treatment duration was 45 days. MWM: Morris water maze, HIP: hippocampus. **B)** <u>Left</u>: Golgi-Cox stained CA1 area with adjacent neocortex below; scale bar: 200 µm. <u>Right</u>: yellow box in stratum radiatum on the left shown at higher magnification. Blue Methoxy-X04 amyloid-β plaque encircled by microglia and reactive astrocytes. Yellow lines indicate the distance of the beginning, middle and end point of a spine-counted segment of a secondary apical CA1 pyramidal cell dendrite to determine the mean distance to the closest plaque; scale bar: 10 µm.

Spine densities were assessed by manual counting in Golgi-Cox-stained brain sections. Secondary dendrites of the stratum radiatum of hippocampal CA1 pyramidal neurons were analyzed. In APP/PS1 mice, Aβ plaques were visualized with blue fluorescent methoxy X04. Since reduction in spine numbers in APP/PS1 mice strongly depends on proximity to Aβ plaques (Kartalou *et al*., 2020a; Kartalou *et al*., 2020b), dendrites were categorized into two groups: secondary dendrites <50 µm away from the closest plaque were termed “near”, whereas dendrites 50-100 µm away were termed “distant” dendrites (compare **Fig.2**).

**Figure 2.**
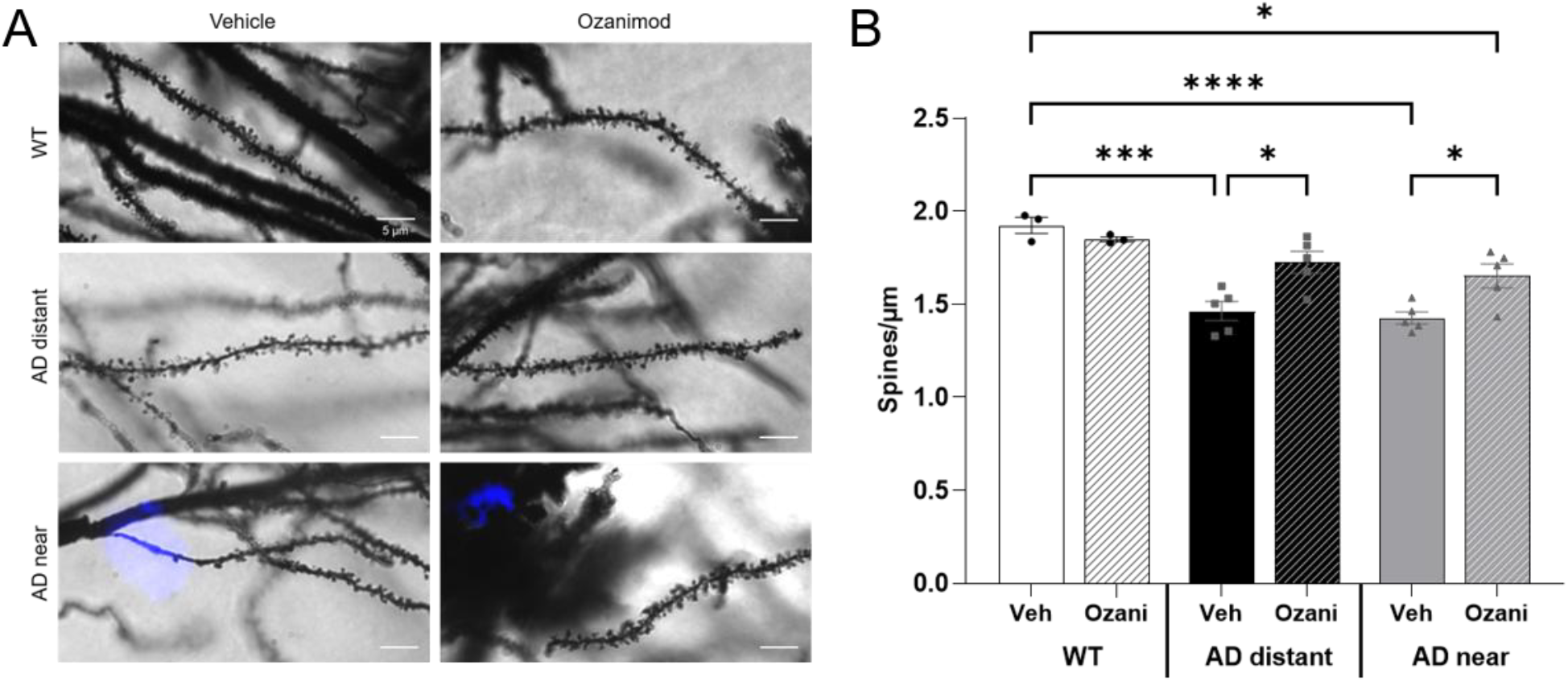
Treatment with Ozanimod mitigates reduced spine densities in dendrites of CA1 pyramidal neurons of APP/PS1 mice. **A)** Spine densities in secondary apical dendrites of hippocampal CA1 pyramidal cells were analyzed in 16months old APP/PS1 mice, after 4-6 weeks treatment with ozanimod via drinking water. The photographs show typical results for different groups as indicated scale bars: 5 µm). **B)** Analysis of dendritic segments: “distant” = 50-100 µm, and “near” = <50 µm away from closest Aβ plaque. Ozanimod significantly inhibited spine density decline compared to vehicle treated controls in dendritic segments both near and distant to Aβ plaques. No effect of ozanimod was observed in WT animals. Data are presented as mean ± SEM; n = 30-50 dendrites from 3-5 mice per group were analyzed. Statistical significance was set at P < 0.05. * = P < 0.01, ** = P < 0.001, *** = P < 0.0001 (post-hoc Tukey comparisons).

We observed a significantly higher spine density in dendrites from the WT vehicle group compared to distant and near dendrites of vehicle treated APP/PS1 mice (**Fig.2B;** WT + vehicle: 1.92 ± 0.04 spines/µm dendrite; APP/PS1 + vehicle near: 1.43 ± 0.03; APP/PS1 + vehicle distant: 1.46 ± 0.05). Importantly, Ozanimod treatment resulted in complete rescue of spine deficits in dendrites near and distant to Aβ plaques in APP/PS1 + RPC1063 mice compared to WT + RPC1063 (APP/PS1 + RPC1063 near: 1.65 ± 0.06 APP/PS1 + RPC1063 distant: 1.73 ± 0.06; WT + RPC1063: 1.85 ± 0.01). Moreover, the rescued spine densities in Ozanimod treated APP/PS1 mice did not differ significantly from vehicle-treated WT animals. However, spine densities in near dendrites from treated APP/PS1 mice could not be completely rescued compared to vehicle-treated WT animals. This indicates that the already existing decline in spine densities that is observed in untreated 6-7 months old APP/PS1 animals (compare Kartalou et al., 2020a; 2020b), which likely worsened until 16 months of age, is partially reversed to levels of age-matched untreated WT animals. Overall, a two-way ANOVA analysis revealed significant main effects for the factors genotype and treatment (F’s ≥ x.y, p’s ≤ 0.034), as well as a significant interaction of these two factors (genotype x treatment: (F2, 174 = 21.51, p<0.0001)).

Interestingly, a similar full reversion of spine deficits in both near and distant dendrites was observed in 16-17 months old APP/PS1 mice treated with Fingolimod (FTY720; 1 mg/kg bodyweight, see **Supplementary Fig.1**), demonstrating that both S1PR modulators exert comparable rescuing effects even when treatment is initiated substantially late at 10 months after cognitive symptoms onset.

### 2. Effects of Ozanimod on microglia-mediated neuroinflammation

We next examined whether the striking improvement in spine density observed in Ozanimod treated 16-17 months old APP/PS1 mice is accompanied by reduced neuroinflammation in the hippocampus and neocortex. Aβ plaque formation in AD mice triggers microgliosis leading to altered microglia morphology and increased expression of the microglia marker protein Iba1 (compare e.g. Aytan et al., 2016; Kartalou et al., 2020). After 45 days treatment of 16 months old APP/PS1 mice and wt littermate controls with 1 mg/kg/day Ozanimod via drinking water, hippocampal and neocortical sections were prepared for immunofluorescent staining against Iba1 (**Fig.3**). We quantified changes in the percentage area occupied by Iba1 positive microglia. To assess potential additional changes in Iba1 expression, we analyzed the normalized Iba1 intensity, defined as the integrated density of the staining divided by the analyzed section area (**Fig. 3C, D**).

**Figure 3.**
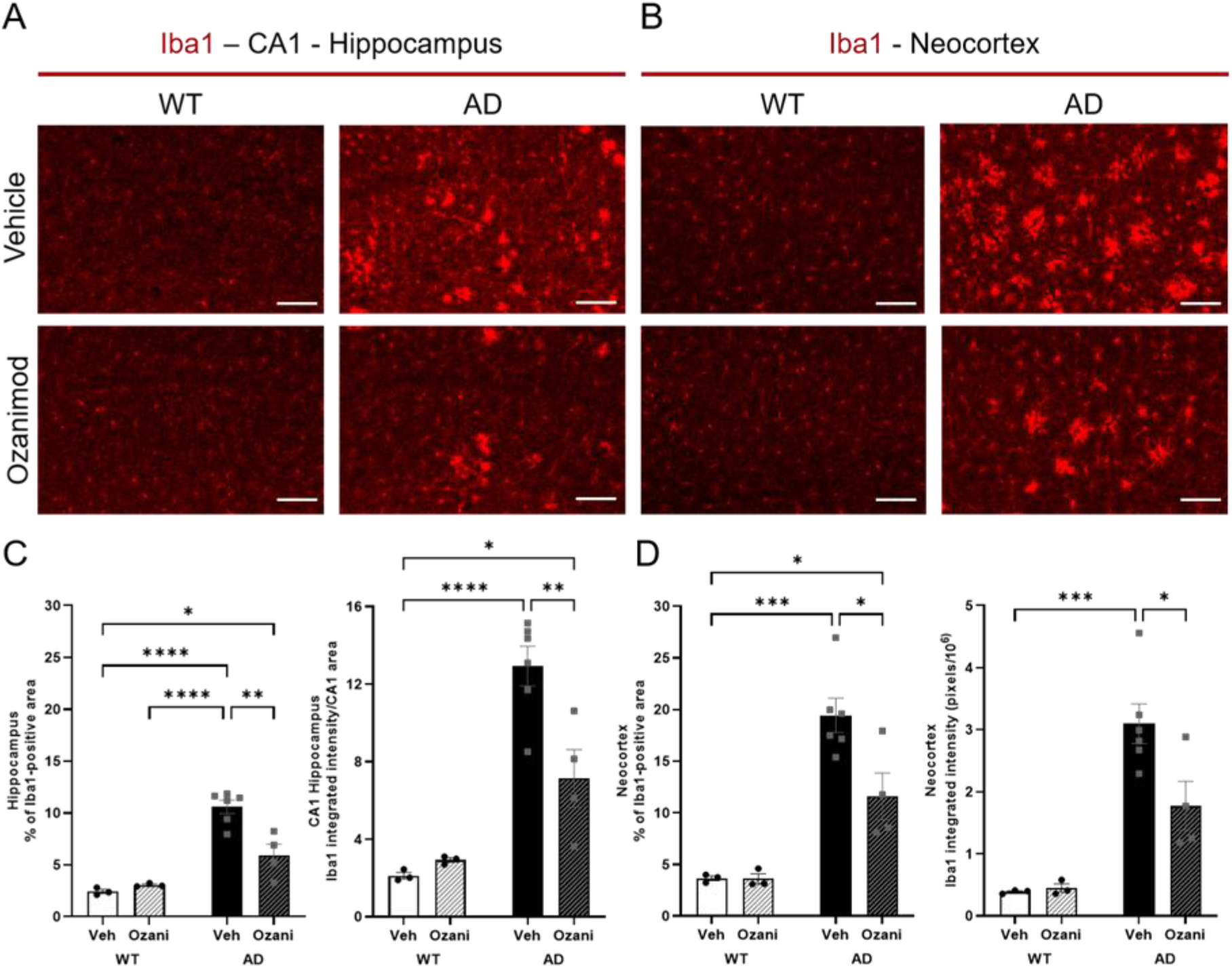
Ozanimod treatment reduces microgliosis in hippocampus and neocortex of 16-17 months old APP/PS1 mice. **A, B)** Iba1 immunofluorescence in hippocampal and neocortical sections of 16-17 months old APP/PS1 mice and wt littermates after 30-45 days treatment with ozanimod (1 mg/kg/day via drinking water). Scale bars: 100 µm. **C, D)** Results for percentage of Iba1 positive area and for integrated intensity per analyzed area of hippocampal and neocortical sections in treated vs. untreated APP/PS1 mice and littermate controls. Ozanimod significantly reduced the increased Iba1 positive area and the integrated intensity per analyzed area that was observed in untreated (vehicle) APP/PS1 mice in hippocampus **(C)** and neocortex **(D)**. No effect of ozanimod was observed in WT animals. Data are presented as mean ± SEM; n = 4 to 6 mice per group with 3 analyzed brain slices per mouse. Statistical analysis was performed using two-way ANOVA followed by a Tukey’s post hoc test. (* = P < 0.05, ** = P < 0.01, *** = P < 0.001. Statistical significance was set at P < 0.05. * = P < 0.01, ** = P < 0.001, *** = P < 0.0001 (post-hoc Tukey comparisons).

We observed a significantly increased percentage of Iba1 positive area and of normalized Iba1 intensity in vehicle treated APP/PS1 mice vs. vehicle treated wt littermate controls in the hippocampus (**Fig. 3C;** HIP area: APP/PS1 + vehicle: 10.58 ± 0.64%; WT + vehicle: 2.43 ± 0.20%; HIP integrated intensity/area: APP/PS1 + vehicle: 12.93 ± 1.02; WT + vehicle: 2.12 ± 0.17). Similar effects were observed in the neocortex (**Fig. 3D**; Neocortex area: APP/PS1 + vehicle: 19.43 ± 1.66%; WT + vehicle: 3.66 ± 0.23%; Neocortex integrated intensity/area: APP/PS1 + vehicle: 3.10 ± 0.32; WT + vehicle: 0.45 ± 0.07).

Importantly, ozanimod treated APP/PS1 mice showed a significant reduction of Iba1 area and of normalized intensity in hippocampus and neocortex compared to vehicle treated APP/PS1 mice (**Fig. 3C;** HIP area: APP/PS1 + RPC1063: 5.91 ± 1.07%; WT + RPC1063: 3.04 ± 0.10%; HIP integrated intensity/area: APP/PS1 + RPC1063: 7.13 ± 1.48; WT + RPC1063: 2.93 ± 0.12; **Fig. 3D**; Neocortex area: APP/PS1 + RPC1063: 11.59 ± 2.27%; WT + RPC1063: 3.61 ± 0.49%; Neocortex integrated intensity/area: APP/PS1 + RPC1063: 1.77 ± 0.39; WT + RPC1063: 0.45 ± 0.07). These results are consistent with reduced microgliosis in hippocampus and neocortex from APP/PS1 mice in response to treatment with Ozanimod.

### 3. Effects of Ozanimod on reactive astrocytes in APP/PS1 mice

Aβ plaque formation and microgliosis are known to induce the transition of resting astrocytes into a reactive state, called reactive astrocytes, characterized by increased surface area and higher expression levels of the astrocyte marker glial fibrillary acidic protein (GFAP) (see e.g. (Heneka *et al*., 2015; Liddelow *et al*., 2017; Liddelow & Sofroniew, 2019; Habib *et al*., 2020)). Therefore, we used immunofluorescent staining against GFAP (**Fig.4**) to quantify the percentage area occupied by GFAP-positive astrocytes in hippocampal and neocortical sections. In addition, we quantified the normalized GFAP fluorescence intensity to detect increased GFAP expression (**Fig.4C, D**). The sections analyzed were adjacent to the respective sections probed for Iba1 immuno-fluorescence and originated from the same mice used to quantify spine densities. Experimental procedures for 16-17 months old animals were identical as described for Iba1 analysis of microgliosis.

**Figure 4.**
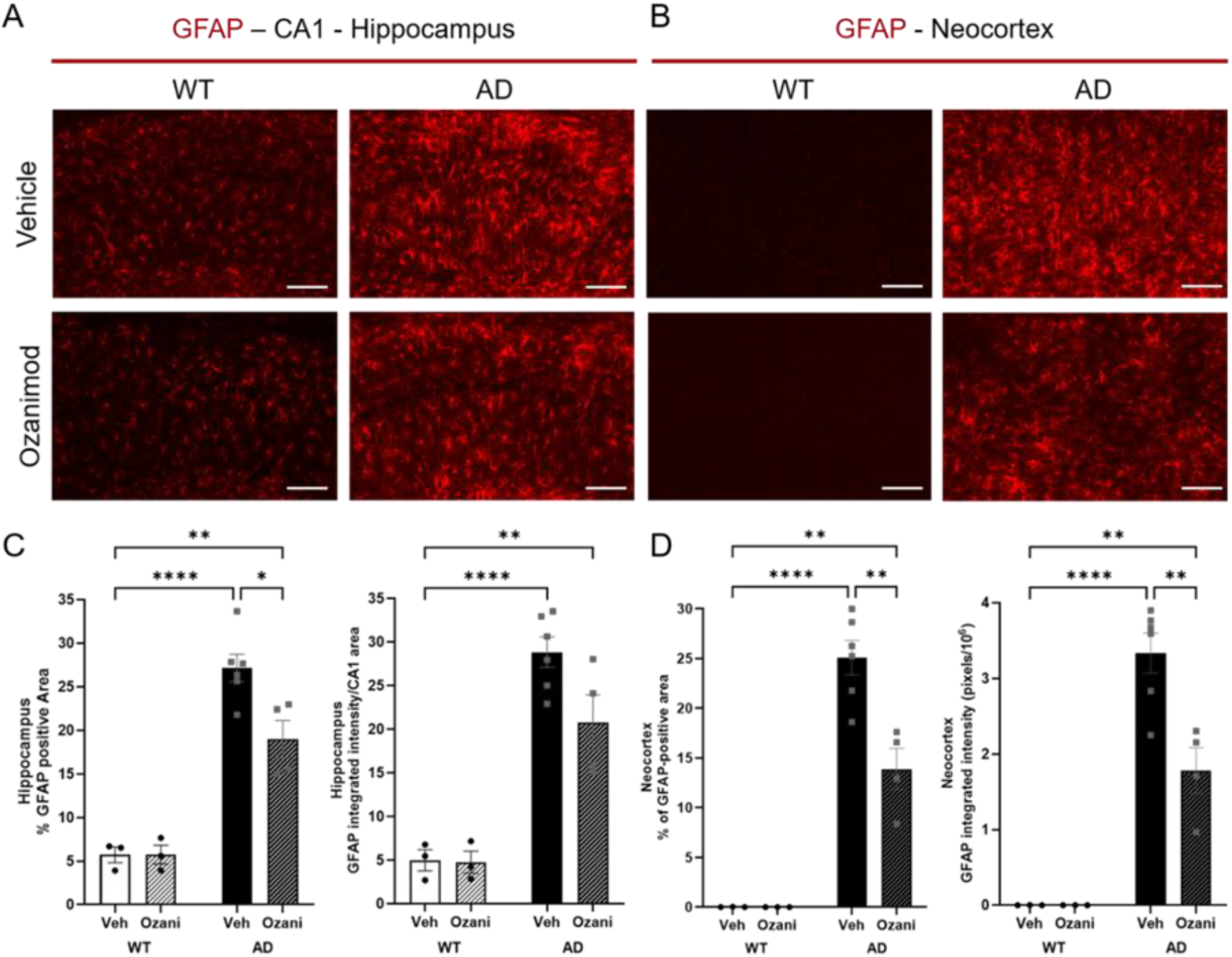
Ozanimod treatment reduces astrogliosis in hippocampus and neocortex of 16-17 months old APP/PS1 mice. **A,B)** GFAP immunofluorescence in hippocampal and neocortical sections of 16-17 months old APP/PS1 mice and wt littermates after 45 days treatment with ozanimod (1 mg/kg/day via drinking water; scale bars: 100 µm). **C, D)** Results for percentage of GFAP positive area and normalized GFAP fluorescence intensity of analyzed hippocampal (C) and neocortical sections (D) in treated vs. untreated APP/PS1 mice and littermate controls. Ozanimod significantly reduced the increased GFAP positive area that was observed in untreated (vehicle) APP/PS1 mice. Ozanimod significantly reduced normalized GFAP fluorescence compared to untreated (vehicle) APP/PS1 mice. No effect of ozanimod was observed in WT animals. The data are presented as mean ± SEM; n = 4 to 6 mice per group, with 3 brain sections analyzed per mouse. Statistical analysis was performed using two-way ANOVA followed by a Tukey’s post hoc test. Statistical significance was set at P < 0.05. * = P < 0.01, ** = P < 0.001, *** = P < 0.0001.

We observed a significant increase in GFAP-positive area and normalized integrated intensity in the hippocampal CA1 subfield in vehicle-treated APP/PS1 mice compared to vehicle wt controls (**Fig. 4C;** HIP area: APP/PS1 + vehicle: 27.16 ± 1.58%; WT + vehicle: 5.71 ± 0.90%; HIP integrated intensity/area: APP/PS1 + vehicle: 28.83 ± 1.76; WT + vehicle: 4.98 ± 1.20). In neocortex of WT animals, GFAP immunoreactivity was below detection, whereas vehicle treated APP/PS1 mice displayed strong GFAP signals (**Fig. 4D**; Neocortex area: APP/PS1 + vehicle: 25.09 ± 1.74%; Neocortex integrated intensity/area: APP/PS1 + vehicle: 3.33 ± 0.27).

Ozanimod-treated mice showed a significant decrease in GFAP-positive area and normalized integrated intensity in the hippocampal CA1 subfield compared to vehicle controls (**Fig. 4C**; HIP area: APP/PS1 + RPC1063: 18.98 ± 2.15%; WT + RPC1063: 5.74 ± 1.08%; HIP integrated intensity/area: APP/PS1 + RPC1063: 20.77 ± 3.17; WT + RPC1063: 4.74 ± 1.27). In the neocortex above the CA1 area, GFAP values were also significantly decreased compared to vehicle-treated APP/PS1 controls (**Fig.4 D**; Neocortex area: APP/PS1 + RPC1063: 13.88 ± 2.09%; Neocortex integrated intensity/area: APP/PS1 + RPC1063: 1.78 ± 0.30). Overall, these results revealed a significant reduction of reactive astrocytes in hippocampus and neocortex of Ozanimod-treated APP/PS1 mice.

### 4. Effects of Ozanimod on Aβ-pathology in APP/PS1 mice

In this series of experiments, we asked whether Ozanimod treatment, in addition to the reduction of microgliosis and astrocytosis, also affects Amyloid-β plaque load. We used Thioflavine S staining to assess changes in number and size of Aβ plaques in the hippocampus and neocortex of 16-17 months old APP/PS1 mice treated with Ozanimod for 45 days (**Fig. 5A**; adjacent slices of animals also analyzed for Iba1 and GFAP; same mouse cohort as for spine density analysis). In the hippocampal CA1 region, Ozanimod treatment significantly reduced the percentage area covered by plaques to roughly 60% (AD vehicle: 2.69 ± 0.21%, AD + RPC1063: 1.64 ± 0.38; **Fig. 5C**). A similar significant decrease was observed for the number of Aβ plaques per unit area upon ozanimod treatment (AD vehicle: 198.07 ± 17.42 plaques/mm^2^; AD + RPC1063: 124.24 ± 27.94), whereas the average Aβ plaques size was not affected (AD vehicle: 140.69 ± 7.38 µm^2^; AD + RPC1063: 133.12 ± 10.68). In the neocortex, plaque burden was significantly reduced to roughly 65% compared to untreated controls (**Fig. 5C;** AD vehicle: 5.00 ± 0.47% ; AD + RPC1063: 3.27 ± 0.45), whereas plaque number per unit area (AD vehicle: 344.78 ± 37.60 plaques/mm^2^; AD + RPC1063: 247.92 ± 43.03) and plaque size (AD vehicle: 149.76 ± 9.25 µm^2^; AD + RPC1063: 136.64 ± 3.50) remained unaffected. Overall, this analysis revealed a significant reduction in Aβ plaque load in hippocampus and neocortex, as well as in plaque number in the neocortex, but a modest decrease in plaque size and in plaque number in the hippocampus in response to ozanimod treatment.

**Figure 5.**
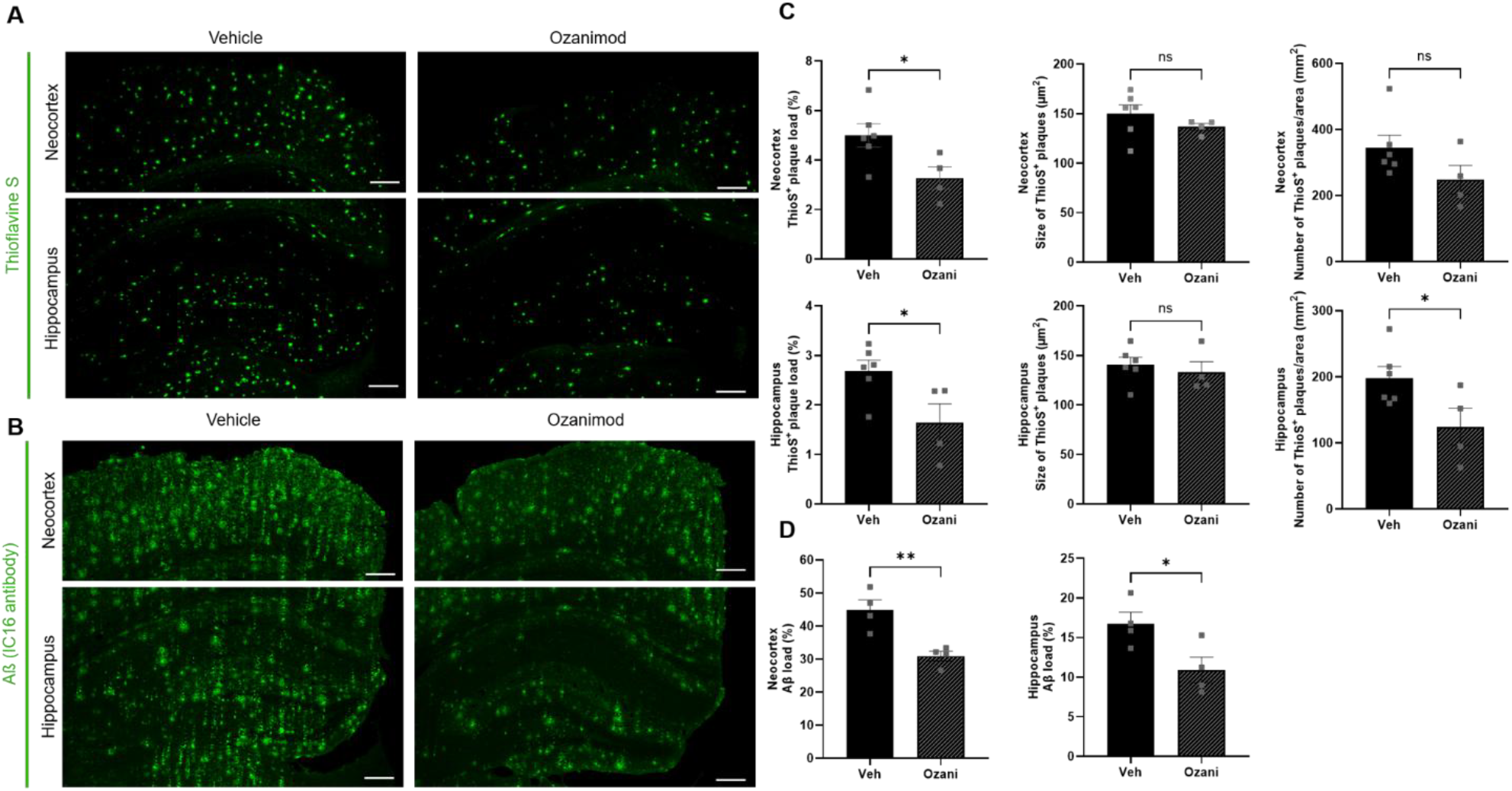
Ozanimod reduces Aβ-plaques and overall Aβ-load in hippocampus and neocortex of 16-17 months old APP/PS1 mice. **A)** Aβ plaques in hippocampus and neocortex were stained with green fluorescent Thioflavine S in sections of 16-17 months old APP/PS1 mice and wt littermates after 45 days of ozanimod treatment (1 mg/kg/day via drinking water). **B)** Overall Aβ protein deposits were detected with an anti-amyloid-β antibody and labeled with a green fluorescent secondary antibody in hippocampal and in neocortical sections. Illumination settings and exposure times as well as thresholding procedures of pictures for quantitative analysis were identical for sections from vehicle and ozanimod treated animals, respectively, for both types of staining. Scale bar: 200 µm in all images. **C)** In hippocampus, ozanimod significantly reduced the percent area and the number of plaques compared to vehicle control, whereas in neocortex, only the percent area was significantly reduced. Average plaque size was neither affected in hippocampus nor in neocortex. **D)** Quantification of anti-amyloid-β-load, expressed as the percentage area of analysed sections. Aβ immunoreactivity in hippocampus and neocortex was significantly reduced in Ozanimod treated APP/PS1 mice compared to vehicle controls. Horizontal bars indicate statistical significance between selected groups. Significance of differences was tested with unpaired Student’s t-test (n=6 animals per group). Significance level was set to 0.05 (p<0.05). Different levels of significance are indicated by stars, with * = p<0.05, ** p< 0.01 (post-hoc Tukey comparisons).

We also performed anti-Aβ immunohistochemical staining, which detects, in addition to Aβ plaques, also less condensed Aβ protein deposits, including oligomers and fibrils (**Fig. 5A**). As expected, this analysis yielded substantially higher values for the percent area covered by Aβ positive material than the Thioflavine S staining (hippocampus:16.74 ± 1.46%, neocortex: 44.90 ± 3.00%; **Fig.5D**) in vehicle treated APP/PS1 mice. Following treatment with ozanimod, a slight reduction to roughly 70% of untreated controls was observed (hippocampus + RPC1063: 10.92 ± 1.60%, neocortex + RPC1063: 30.62 ± 1.47%).

These data indicate that Ozanimod significantly reduces Aβ burden to a similar extent as microgliosis and astrocytosis.

## Supplementary Figure

**Supplementary Fig. S1.**
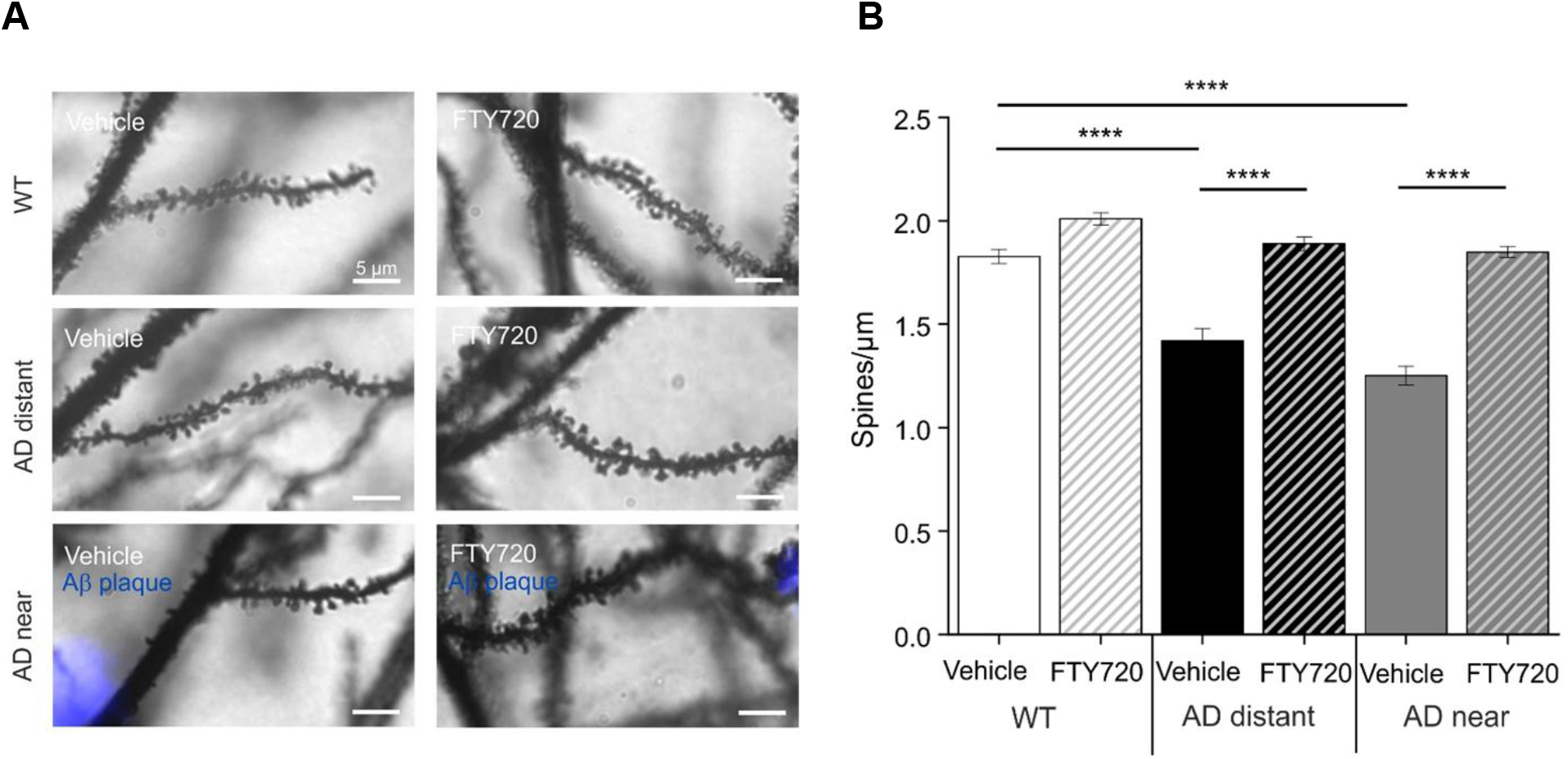
Fingolimod reduces spine deficits in CA1 of hippocampus in 15 months old APP/PS1 mice. **A)** Male mice were treated for 4 weeks with Fingolimod (1 mg/kg body weight). Secondary apical dendrites of CA1 pyramidal neurons from 15 months old mice in vehicle and fingolimod treated wild type (WT) animals (WT, upper panel), in APP/PS1 animals >50 μm away from the closest plaque (AD distant, middle panel), or in APP/PS1 mice <50 μm away from the nearest plaque (AD near, lower panel). Scale bar: 5 μm in all images. **B)** Quantification of spine densities in secondary apical CA1 dendrites shown for the six different groups in A. Each bar represents the mean value of 10 dendrites from 10 different CA1 pyramidal neurons per animal (three animals per group). Horizontal bars with asterisks indicate statistical significance between selected groups. Significance of differences was tested with two-way ANOVA followed by Tukey’s post hoc test (n = 30 dendrites from three animals per group). Significance level was set to 0.05 (p < 0.05). Levels of significance are indicated by asterisks, with **** = p < 0.0001 (post-hoc Tukey comparisons).

## Discussion

To the best of our knowledge, our current investigation is the first study to report effects of the S1PR modulator Ozanimod on AD pathology in a mouse model. Ozanimod effectively attenuated dendritic spine loss in hippocampal CA1 neurons in our APP/PS1 mouse model, even when treatment was initiated at 16 months of age, after cognitive AD symptoms had already persisted for 10 months. Ozanimod treatment also reduced microgliosis, astrogliosis, and Aβ burden in hippocampus and neocortex of 16-17 months old APP/PS1 mice after 45 days of treatment. Because Ozanimod is already an FDA and EMA approved drug for treatment of multiple sclerosis, our findings provide a rationale for further clinical trials, testing ozanimod as a potential anti-neuroinflammatory therapy in human AD patients.

The anti-MS drug Fingolimod is a structural analog of endogenous sphingosine-1-phosphate and the founding member of a family of sphingosine-1-phosphate receptor (S1PR) modulators that have recently shown potential to be effective also against other (neuro-)inflammatory diseases (see e.g. (Angelopoulou & Piperi, 2019; Bascunana *et al*., 2020; Pournajaf *et al*., 2022)). Fingolimod was shown previously to be protective against pathology in an AD mouse model (Aytan *et al*., 2016; Carreras *et al*., 2019). Of note, more recent studies suggested that Fingolimod counteracts or even reverses cognitive decline and synaptic dysfunctions in different AD mouse models when treatment started at the stage of emerging memory impairment (Kartalou *et al*., 2020b; Baloni *et al*., 2022). Among the other S1PR modulators, only Siponimod has recently been tested in an AD mouse model and showed beneficial effects against Aβ induced pathology (Zhu *et al*., 2023).

### Amelioration of spine pathology in APP/PS1 mice by Ozanimod

We showed previously that mice of this APP/PS1 model (Radde *et al*., 2006) develop spine loss in hippocampal CA1 neurons in parallel with reduced long-term potentiation (LTP) at glutamatergic inputs to CA1 and deficits in hippocampus-dependent spatial memory formation (Kartalou *et al*., 2020b). In the same study, we observed that 4-5 weeks of Fingolimod treatment of 5-6 months old APP/PS1 mice partially reversed these synaptic and memory deficits. In the present study, we observed that oral Ozanimod application via drinking water, starting 10 months after persisting learning and memory deficits in these mice (at 16 months of age; **Fig. 1A**), counteracted spine pathology to a similar extent as we previously observed with i.p. injections of Fingolimod at younger age in these APP/PS1 mice (Kartalou *et al*., 2020b). We thus conclude that Fingolimod and Ozanimod are comparably effective to ameliorate spine deficits in our APP/PS1 mouse model. This conclusion is supported by our observation that Fingolimod treatment starting at 16 months of age reduced spine pathology to a similar extent as Ozanimod (compare **Supplementary Fig. S1**). This is of interest because Ozanimod and Fingolimod show distinct binding efficacies to the 5 different S1PR subtypes (S1PR1-5). Ozanimod and its main metabolites bind *in vivo* to S1PR1 and S1PR5, with a 10-fold preference for S1PR1. In contrast, Ozanimod does not show significant binding to S1PR2, S1PR3, and S1PR4 receptors. In MS, Ozanimod and Fingolimod exert their main effect (reduced egress of lymphocytes from lymph nodes) through functional antagonism at S1PR1, whereas it is still unclear whether Ozanimod exerts a relevant effect on brain parenchymal cells via S1P5. Fingolimod shows comparable potency at S1PR1 as Ozanimod, but a much higher affinity for binding to S1PR5 than Ozanimod. Moreover, Fingolimod – but not Ozanimod – additionally binds to S1PR3 and S1PR4 (reviewed in (Chaudhry *et al*., 2017); (Angelopoulou & Piperi, 2019). Taken together, these data suggest that the shared spine rescuing effect of Ozanimod and Fingolimod is mediated primarily by S1PR1. However, additional studies are needed to identify the S1P receptor subtypes that inhibit the synaptic pathology in AD mouse models.

### Amelioration of microgliosis and astrocytosis in APP/PS1 mice by oral administration of Ozanimod

Our results for Iba1-positive microglia and GFAP-positive astroglia in this investigation are consistent with many previous studies obtained with Fingolimod in different AD mouse models (e.g. (Aytan *et al*., 2016; Carreras *et al*., 2019; Kartalou *et al*., 2020b; Baloni *et al*., 2022); reviewed in (Lessmann *et al*., 2023)). These previous studies described that Fingolimod i.p. injections or application via drinking water in 6-8 months old animals from different AD mouse models reduces microglia induced neuroinflammation. Our current study extends these findings by showing that oral treatment with Ozanimod also in 16-17 months old mice significantly reduced Iba1 staining of microglia in hippocampus and neocortex of APP/PS1 mice (**Fig.3**). This is an important finding since it shows that much delayed treatment of AD mice starting 10 months after fully developed memory deficits S1PR modulators can still reverse long established synaptic dysfunctions. The Iba1 staining reduction in neocortex of 16-17 old animals by Ozanimod to roughly 50% of vehicle control was similar to our previous results with Fingolimod in 6-7 months old mice (Kartalou *et al*., 2020b). However, the reduction of Iba1 positive microglia with Ozanimod in hippocampus was less efficient than observed with Fingolimod in these younger animals.

Similar to observations for Iba1, also the reduction of percent area and integrated intensity per unit area of GFAP positive astrocytes in response to Ozanimod treatment in 16-17 months old APP/PS1 mice was less pronounced in hippocampus and neocortex (**Fig.4**) than previously described after Fingolimod treatment (Kartalou *et al*., 2020b). Given that not many previous studies in AD mouse models investigated 16-17 months old animals, it is conceivable that neuroinflammatory processes in AD mice of this age might have reached a chronic stage that cannot be completely reversed by 45 days of Ozanimod treatment. Interestingly, the similar efficacy of Fingolimod in previous studies and Ozanimod in the present investigation to reduce microglia and astroglia pathology speaks in favour of mutual dependence of both types of cells in neuroinflammation. This is consistent with the view that activated microglia induce the formation of reactive astrocytes and both cell types orchestrate neuroinflammation in AD and other neurodegenerative diseases (Liddelow *et al*., 2017; Joshi *et al*., 2019; Liddelow & Sofroniew, 2019). Whether longer treatment periods or different treatment regimens with S1PR modulators might more effectively restore activated microglia and astrocytes to a homeostatic state remains to be determined.

### Reduction of Aβ plaque load and Aβ immunolabelling in APP/PS1 mice by oral Ozanimod treatment

Fingolimod induced reduction of Aβ pathology and downstream effects on neurodegeneration and memory were among the first outcomes that were reported for S1PR modulators in AD and other neurodegenerative diseases models (Asle-Rousta *et al*., 2013; Doi *et al*., 2013; Takasugi *et al*., 2013; Fukumoto *et al*., 2014; Miguez *et al*., 2015). Moreover, Fingolimod was reported to reduce Aβ plaque burden in many different transgenic AD mouse models (reviewed e.g. in (Angelopoulou & Piperi, 2019; Bascunana *et al*., 2020; Baloni *et al*., 2022; Lessmann *et al*., 2023). Likewise, we also observed in our current study a reduction in Aβ covered area in hippocampus and neocortex in response to Ozanimod treatment of APP/PS1 mice (**Fig.5**). On the same vein, mono- and oligomeric Aβ species outside of Aβ plaques were significantly reduced, as was evident from our Aβ immunolabeling with the IC16 antibody. More recently, another S1PR modulator, Siponimod, was shown to exhibit beneficial effects in a transgenic AD mouse model (Zhu *et al*., 2023). Together, these findings suggest either a general inhibitory effect of S1PR1 modulators on Aβ plaque formation or a positive effect on Aβ plaque clearance. Whether these Aβ reducing effects are mediated by S1PR signaling in microglia and/or astroglia, or whether S1PR-mediated Aβ reduction occurs upstream from microglia and astrocyte signaling, thereby reducing the amount of activated microglia and reactive astrocytes, remains to be investigated.

## Conclusion

One of the main objectives of AD research is to identify therapeutic approaches that effectively counteract cognitive decline. The results of the present study suggest that oral application of the anti-inflammatory S1PR1- and S1PR5-specific modulator Ozanimod (RPC1063) via drinking water can reverse synaptic spine deficits in the hippocampus, as well as microgliosis, astrocytosis, Aβ plaque- and Aβ oligomer load in hippocampus and neocortex of an APP/PS1 AD mouse model, even when treatment starts at 16 months of age, when AD symptoms are manifested for already 10 months. In conjunction with numerous previous preclinical studies demonstrating a similar AD-rescuing effect of the related unspecific S1PR modulator Fingolimod (FTY720), our findings suggest that anti-inflammatory treatment with S1PR-modulators might be beneficial for treatment of AD patients either with Ozanimod/Fingolimod monotherapy, or as co-therapy with monoclonal Aβ antibodies.

## Materials and Methods

### Animals

For experiments, 16-17 months old heterozygous APP/PS1 mice and their wildtype littermates were used. These APP/PS1 mice harbor a mutation in the APP gene (KM670/671NL, “Swedish mutation”) as well as a mutated presenilin 1 (Leu166Pro mutation), both under control of the Thy1 promotor that drives neuron specific expression of these proteins [17]. Both gene mutations are associated with early-onset familial AD (FAD) in humans. Experimental procedures in this investigation were similar as described previously in Kartalou et al. (2020b), with minor modifications. The animals were generated on a C57BL/6J genetic background and constantly backcrossed with C57BL/6J mice (Charles River, Sulzfeld, Germany). The animals were housed in groups of 3-4 animals and had constant access to food and water. They were maintained on a 12:12 h light dark cycle (lights on at 7 a.m.). All experiments were performed during the light period of the animals, were in accordance with the European Committees Council Directive (2010/63/EU) and approved by the local animal care committees.

### Ozanimod (RPC1063) administration

Male WT and AD transgenic mice were treated via drinking water with Ozanimod (RPC1063) at a concentration of 1 mg/kg body weight for 45 days. To ensure accurate dosing, average daily water consumption per cage was monitored, and the Ozanimod concentration in the drinking water was adjusted accordingly to achieve the intended dose of 1 mg/kg body weight per animal. Control mice were treated identically with vehicle solution. All animals were treated according to this scheme until they were sacrificed for dendritic spine quantification or immunohistochemical experiments. Spine analysis was conducted in the same cohort of animals as the immunohistochemical stainings. Right brain hemispheres were used for spine density analysis and left hemi-brains for GFAP, Iba1, ThioS and Abeta analyses.

### Combined Aβ plaque staining and Golgi-Cox impregnation

All mice were injected twice (24h interval) i.p. with 75 µl of 10 mg/ml Methoxy-X04 (TOCRIS) in DMSO (Sigma-Aldrich) as described previously (Jahrling *et al*., 2015; Bayram-Weston *et al*., 2016). Two hours after the second injection, they were anesthetized and transcardially perfused with 0.9% saline followed by 4% PFA/PB (pH 7.4). The injection steps were performed according to Jahrling et al. (2015). The brains were divided into two hemispheres. Both hemispheres were postfixed in 4% PFA/PBS pH,7.4 for 24 hours at 4 °C. The left brain hemispheres were prepared for immunohistochemical stainings and the right hemi-brains were transferred into Golgi-Cox solution in the dark and the solution was changed only once after 7 days. After 14 days, the hemi-brains were placed into 25% sucrose in PBS at 4 °C for at least 1-2 days. Coronal sections of 100 μm thickness were cut using a Vibratome (Pelco Model 1000, The Vibratome Company, St. Louis, USA). Sections were mounted onto gelatine-coated slides and after allowing them to dry, they were washed with distilled water for 2 min and then they were transferred into 20% ammonium hydroxide in distilled water (Ammonium hydroxide solution, ACS reagent, 28.0 – 30.0 % NH3 basis, Sigma-Aldrich). The sections were washed again with distilled water twice for 2 min each. The sections were dehydrated passing through ascending grades of ethanol 70%, 95% and 100% for 5 min each and cleaned in Xylol (Roth) twice for 10 min each. The sections were coverslipped with DePex medium (Serva) and after letting them dry for 2 days they stored at 4 °C ready for analysis. The composition for the Golgi-Cox solution was 5% potassium dichromate (Merck), 5% mercuric chloride (Merck) and 5% potassium chromate (Merck; compare [41]). All stock solutions as well as the final Golgi-Cox solution were prepared according to the protocol used by Bayram-Weston (Bayram-Weston *et al*., 2016).

### Image processing and dendritic spine density analysis

Spine density of secondary apical dendritic segments in stratum radiatum (SR) of CA1 pyramidal neurons was calculated as the number of spines per micrometer dendritic length. Ten dendritic segments (15 µm long) per animal, from 10 different neurons throughout intermediate and ventral hippocampus, were analyzed for all treated and untreated groups. Dendritic lengths were estimated using the NeuronJ plugin of NIH ImageJ software (https://imagej.nih.gov/ij/). For AD mice, 10 dendritic segments which were at a distance more than 50 μm from the plaque border (AD distant) and 10 dendritic segments which were located within 50 μm from the plaque border (AD near) were analyzed per animal. This digital near/distant classification scheme was selected to facilitate quantification of the results. When plotting spine densities versus plaque distance for individual dendritic segments we did not observe a clear cut-off at a certain distance, but a decrease at distances <50 µm is evident (compare Suppl. Fig. 4). Images for Golgi-Cox stained dendritic segments and the blue fluorescent methoxy-X04 stained Aβ plaques were captured with a 40x magnification objective using a SPOT digital camera that was attached to a LEITZ DM R microscope (Leica). The distance in z-direction of analyzed dendrites to the border of the Methoxy-X04 stained Aβ plaque was in all cases <5 µm. This deviation was minor compared to the calculated distances in the x-y space and was therefore neglected. Imaging for Golgi-Cox staining was performed in bright field mode while Methoxy-X04 fluorescence was captured with a filter cube (excitation filter: BP 360/40 nm, dichroic mirror: 400 nm, emission filter: LP 425; Leica). The distance of the analyzed dendritic segments from the Aβ plaques was determined as the average distance of both segment end-points and the center of the segment to the respective plaque border in merged pictures using ImageJ software. Dendritic segments from different neurons were traced and spine density was quantified as the number of spines per dendritic length in µm. The spines were counted manually using a 100x magnification oil immersion objective. Spine density analysis was conducted blindly to the treatment in WT and AD mice.

### Preparation of slices for IHC

The remaining left hemi-brains were transferred to 25% sucrose (AppliChem) in PBS for cryoprotection. The fixed hemispheres were then cut to 40 µm thick coronal slices using a cryostat (Leica CM 3050). Three free-floating sections 240 μm apart from each other (between Bregma levels -2.46 mm and -3.08 mm according to Franklin and Paxinos [43]), containing the hippocampus and cortex were selected for all histochemical fluorescent stainings. For imaging, all sections were transferred to slides (Superfrost, Thermo Scientific), coverslipped with ImmunoMount mounting medium (Thermo Scientific).

### Immunofluorescence for total Aβ

Free-floating sections were permeabilized with PBS/0.1 % Triton X-100 and incubated with 98 % formic acid to perform antigen retrieval. This step was followed by washes with PBS. Sections were then blocked in 20 % normal goat serum (Dianova) in PBS/Triton X-100 0.1 % and incubated overnight at 4°C with the mouse anti-Aβ IC16 antibody (1:400; kindly provided by Prof. Claus Pietrzik, Johannes-Gutenberg-University Mainz [44] in 10 % normal goat serum (Dianova) in PBS containing 0.1 %Triton X-100. Sections were then washed with PBS/0.1 %Triton X-100, incubated with an Alexa Fluor 488 labeled goat anti-mouse antibody (1:500, Thermo Scientific) in 10 % normal goat serum (Dianova) in PBS/0.1 % Triton X-100, and finally washed with PBS. Imaging was performed with ZEN 2010 software using a 5x objective in a confocal imaging system (LSM 780, Zeiss, Germany). Green fluorescence was excited using the 488 nm laser line from an Argon laser. For Aβ load quantification 16 bit images were converted to 8 bit gray-scale images and after defining the region of interest (total hippocampal or cortical area) they were thresholded within a linear range using the NIH ImageJ software. The load was expressed as the percentage area covered by Aβ-positive staining (% Aβ load). The quantification performed blind to the treatment for WT and AD mice. The quantification was performed blind to the treatment for WT and AD mice.

### Immunofluorescence for GFAP

For GFAP staining, antigen retrieval was performed with sodium citrate buffer (10 mM, pH 6.0) for 30 min at 80 °C followed by 3 washes with TBS. Sections were blocked in 5% normal goat serum (Dianova) in TBS with 0.4 % Triton X-100 and incubated with a mouse anti-GFAP antibody (1:500, clone G-A-G, Sigma: G 3893) in blocking solution overnight at 4°C. Sections were washed with TBS and incubated with an Alexa Fluor 488 labeled goat anti-mouse antibody (1:500,Thermo Scientific) in blocking solution and were finally washed with TBS. For the quantification of GFAP in the CA1 area of the hippocampus and neocortex images were captured with a 10x objective using a SPOT camera attached to a Leica LEITZ DM R microscope through an appropriate filter cube (excitation filter: BP 515-560 nm, dichroic mirror: 580 nm, emission filter: LP 590 nm, Leica). Using NIH ImageJ software for quantification, the GFAP load was expressed as the percentage area covered by GFAP-positive staining, and the normalized integrated intensity (raw integrated density divided by the area of the analyzed CA1 area) or the integrated intensity (for identical sized neocortex pictures) were expressed in pixels (pixels/10^6^). The quantification was performed blind to the treatment for WT and AD mice.

### Immunofluorescence for Iba1

Free-floating sections were first treated with sodium citrate buffer (10 mM, pH 6.0) for 30 min at 80°C for antigen retrieval and then washed 3 times with PBS. Sections were blocked in 10 % FBS (Gibco) and 1 % BSA (Sigma) in PBS containing 0.3 % Triton X-100 and incubated overnight at 4°C with a rabbit anti-Iba1 antibody (1:500, Wako) in PBS containing 1 % FBS (Gibco), 0.1 % BSA (Sigma) and 0.3 % Triton X-100. Sections were then washed with PBS and developed with an Alexa Fluor 555 labeled donkey anti-rabbit antibody (1:500, Thermo Scientific) in PBS containing 1 % FBS, 1 % BSA, and 0.3 % Triton X-100. The sections were finally washed with PBS. Imaging and quantification were performed as described for GFAP immunohistochemistry. The quantification was performed blind to the treatment for WT and AD mice.

### Thioflavine S staining

Coronal free-floating sections were incubated for 9 min in 1% Thioflavine S (Sigma-Aldrich) aqueous solution and then treated with 80% ethanol 2 times for 3 min each, followed by a wash with 95% ethanol for 3 min. Sections were rinsed 3 times with distilled water. Thioflavine S imaging was performed with a confocal Zeiss laser-scanning-microscope using Zen software (LSM 780, Zeiss, Germany). A 5x magnification objective was used and green fluorescence was excited with a 488 nm Argon laser. Primary images were converted to 8 bit gray-scale and after defining the region of interest (total hippocampal or cortical area) they were thresholded using NIH ImageJ software. Thioflavine S plaque load was expressed as the percentage area covered by Thioflavine S-positive staining (% ThioS). The ImageJ tool ‘Analyze particles’ was used to determine plaque number and size (plaque area in µm^2^). The quantification was performed blind to the treatment for WT and AD mice.

### Statistical analysis

All data were analyzed by a two-way ANOVA (analysis of variance) using genotype and treatment as between-subject factors. In case of significant main effects, Tukey post-hoc comparisons were performed. A p-value < 0.05 was considered as significant difference. All data were analysed using GraphPad Prism version 8.4 (GraphPad Software, USA).

## Author Contributions

Experiments were performed by: RF (spine analyses, all immunohistochemical procedures, all fluorescence microscopy data); TE (planning of animal treatments); LS (treatment of animals); G-IK (suppl. Fig.1). Experiments were designed by G-IK, TE, RF, KG, and VL. The data were analyzed by RF and TE. The study was designed and supervised by VL, KG, and TE. The manuscript was written by VL, KG, RF, and TE with help from G-IK. VL and KG initiated and conceived the project.

## Acknowledgments

The authors want to thank Dr. Thomas Munsch for help with fluorescence and confocal microscopy, and Birgit Adam, Annika, Ritter, Anja Reupsch, for excellent technical assistance.

